# Emergent epileptiform activity drives spinal sensory circuits to generate ectopic bursting in intraspinal afferent axons after cord injury

**DOI:** 10.1101/2023.07.03.547522

**Authors:** Matthew Bryson, Heidi Kloefkorn, Shaquia Idlett-Ali, Karmarcha Martin, Sandra M. Garraway, Shawn Hochman

**Affiliations:** Emory University School of Medicine Department of Cell Biology (30322); Oregon State University Department of Chemical, Biological, and. Environmental Engineering (97331); University of Colorado School of Medicine

## Abstract

Spinal cord injury (**SCI**) leads to hyperexcitability and dysfunction in spinal sensory processing. As hyperexcitable circuits can become epileptiform elsewhere, we explored whether such activity emerges in spinal sensory circuits in a thoracic SCI contusion model of neuropathic pain. Recordings from spinal sensory axons in multiple below-lesion segmental dorsal roots (**DRs**) demonstrated that SCI facilitated the emergence of spontaneous ectopic burst spiking in afferent axons, which synchronized across multiple adjacent DRs. Burst frequency correlated with behavioral mechanosensitivity. The same bursting events were recruited by afferent stimulation, and timing interactions with ongoing spontaneous bursts revealed that recruitment was limited by a prolonged post-burst refractory period. Ectopic bursting in afferent axons was driven by GABA_A_ receptor activation, presumably via shifting subthreshold GABAergic interneuronal presynaptic axoaxonic inhibitory actions to suprathreshold spiking. Collectively, the emergence of stereotyped bursting circuitry with hypersynchrony, sensory input activation, post-burst refractory period, and reorganization of connectivity represent defining features of epileptiform networks. Indeed, these same features were reproduced in naïve animals with the convulsant 4-aminopyridine (**4-AP**). We conclude that SCI promotes the emergence of epileptiform activity in spinal sensory networks that promotes profound corruption of sensory signaling. This corruption includes downstream actions driven by ectopic afferent bursts that propagate via reentrant central and peripheral projections and GABAergic presynaptic circuit hypoexcitability during the refractory period.

## Introduction

An estimated 80% of patients develop neuropathic pain after spinal cord injury (**SCI**) [1]. Our understanding of the etiology of neuropathic pain, and subsequently, successful treatment options, remains limited [2]. Animal studies on mechanisms of SCI neuropathic pain have focused on spontaneous activity in primary afferents [3, 4] and hyperexcitability in spinal sensory pathways [5-8].

In the dorsal horn (**DH**), primary afferents synapse onto excitatory glutamatergic and inhibitory GABAergic interneurons [9, 10]. Subpopulations of GABAergic interneurons form presynaptic axoaxonic synapses onto intraspinal afferent projections, mediating a critical form of negative feedback control [11-15] (also see[16]) called primary afferent depolarization (**PAD**), recorded experimentally as a dorsal root potential (**DRP**). Though PAD is depolarizing due to high intracellular chloride concentration [17], it is functionally inhibitory via mechanisms including shunting [18]. Excessive PAD can generate suprathreshold **ectopic** action potentials termed dorsal root reflexes (**DRRs**) [19]. These ectopic excitatory spikes have been observed in peripheral injury pain models [20-22] and can propagate bidirectionally: reentrant spikes can synaptically excite central circuits [23, 24] while antidromic spikes propagate to peripheral innervation sites [21, 25]. A unique feature of SCI is that loss of descending systems promotes the emergence of synchronous multisegmental DRPs [26, 27]. Whether suprathreshold ectopic spikes (DRRs) also emerge after SCI-induced hyperexcitability remains unstudied.

SCI-induced hyperexcitability is associated with alterations in voltage-gated sodium [28, 29] and potassium channel function in dorsal root ganglia and DH [30-32], as well as perturbations in DH intracellular chloride homeostasis [5, 33-35]. Circuit hyperexcitability through these mechanisms is prominent in the various epilepsies [36-39], and it is possible that SCI-induced sensory circuit hyperexcitability also recruits epileptiform activity. Consistent with this possibility, the convulsant 4-aminopyridine (**4-AP**) generates epileptiform activity in spinal DH circuits [40]. Typical for neuropathic pain, this activity is insensitive to traditional analgesics but depressed by anticonvulsants, which may also treat SCI neuropathic pain [41].

As epileptiform activity emerges from hyperexcitable circuitry, we sought to determine whether it emerges after SCI. Epileptiform activity would be expected to: 1) consist of episodes of stereotyped bursting activity, 2) exhibit synchrony between normally independent circuits, 3) be triggered by sensory input, 4) exhibit a refractory period, and 5) result from changes in circuit function [40, 42-45]. We sought to identify bursting with these characteristics in the spinal cord.

## Results

### Ectopic afferent bursting associates with spinal cord injury (SCI)

Here, we used the mouse thoracic contusion model of neuropathic pain to study sensory hyperexcitability [46, 47]. The contusion SCI population developed and maintained mechanical hypersensitivity (↓ paw withdrawal reflex threshold) after surgical recovery **(Figure 1A)**. This has been associated with a cognitive component of neuropathic pain in other studies [48, 49]. Sensory circuit hyperexcitability was subsequently assessed in an ex-vivo spinal cord preparation with access to multiple segmental dorsal roots (DRs) for stimulation and recording of primary afferents **(Figure 1B)** [50]. Occasional bouts of spontaneous ectopic bursting in afferent axons were seen in both sham and SCI mice. They appeared both independently in single roots and synchronously across multiple roots **(Figure 1C)**. Individual bursts had similar and stereotyped appearance and duration **(Figure 1D**_**1**_), suggesting common circuit recruitment. However, compared to sham, bursts had larger amplitude and higher frequency in SCI mice **(Figure 1D**_**2**_, **1D**_**3**_). Overall, burst frequency showed moderate correlation with paw mechanosensitivity (↓ paw withdrawal reflex threshold [PWT]; **Figure 1E**)

**Figure 1.**
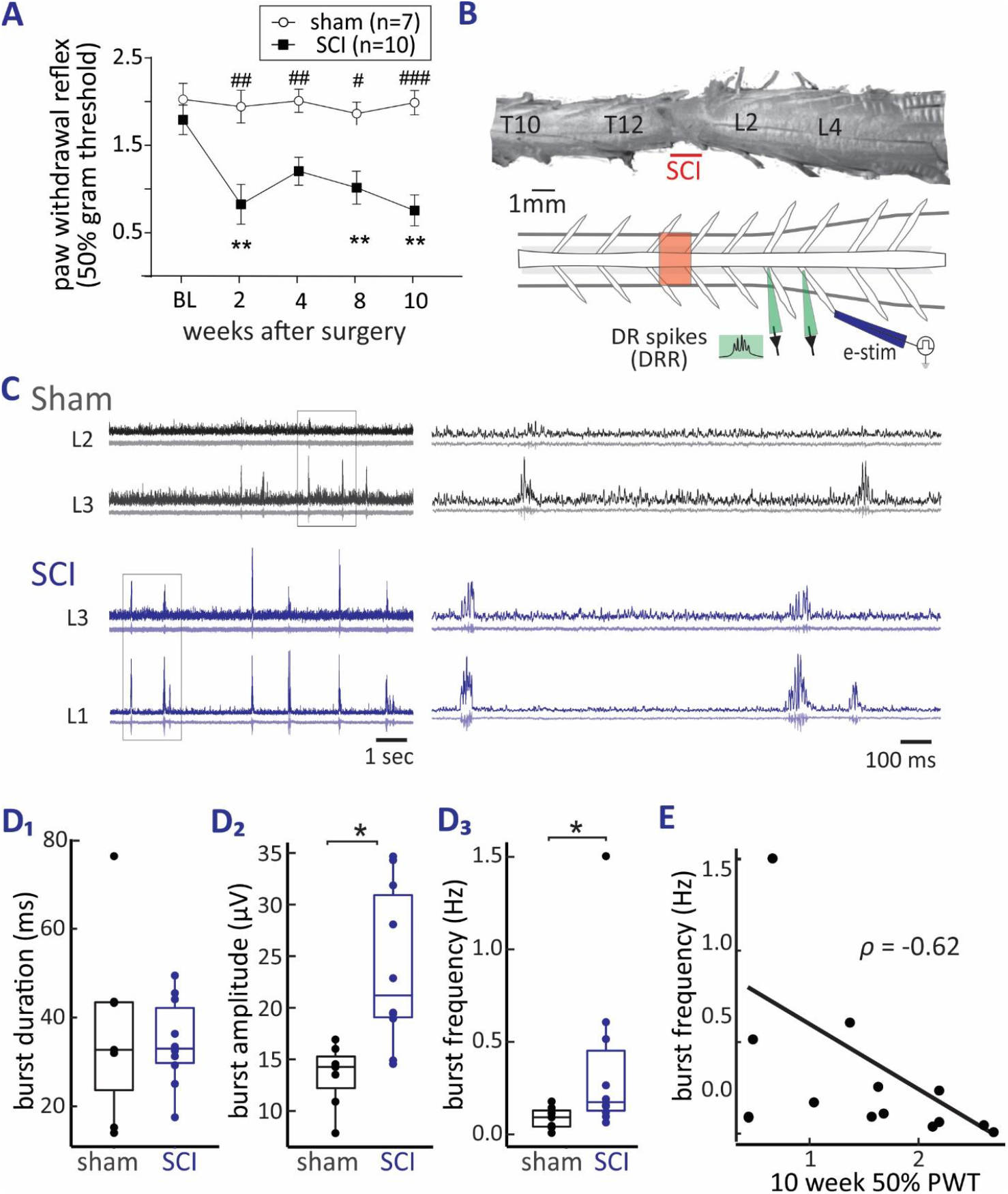
Lower thoracic SCI contusion leads to increased paw mechanosensitivity and promotes spontaneous ectopic bursting in primary afferents. **(A)** SCI but not sham animals developed mechanical hypersensitivity after surgical recovery, as measured by reduction in 50% paw withdrawal threshold. (# compares weekly SCI measure to weekly sham measure; * compares weekly SCI measure to baseline SCI measure; #/*p < 0.05, ##/**p<0.01, ###/***p<0.001) (n = 10 SCI, 7 sham). **(B)** Top: example of SCI spinal cord with labelled roots and injury site. Bottom: diagram of recording configuration. Suction electrodes are attached to dorsal roots and dorsal root entry zones to enable electrical stimulation and recording, respectively. **(C)** Representative spontaneous bursting activity in ipsilateral dorsal roots recorded from an SCI and sham cord preparation. Boxes denote region of magnification in panels at right. Bottom traces for each event are raw recordings. Top traces are RMS amplitude filtered with a 1-3ms time constant. **(D)** Box plots of measured burst parameters in sham and SCI populations. **(D**_**1**_) Burst duration did not differ between sham (44.4 ± 18.2ms) and SCI (35.1 ± 9.5ms) preparations (Wilcoxon rank-sum test p-value = 0.88; n = 10 SCI, 7 sham). **(D**_**2**_) Burst amplitude was higher in SCI (20.6 ± 8.3μV) than sham (15.6 ± 2.2μV) than sham cords (Wilcoxon rank-sum test p = 0.001; n = 10 SCI, 7 sham). **(D**_**3**_) Burst frequency of SCI preparations (0.3 ± 0.4Hz) was greater than that of sham preparations (0.08 ± 0.05Hz). (Wilcoxon rank-sum test p-value = 0.022; n = 10 SCI, 7 sham). **(E)** Paw withdrawal threshold at 10 weeks is negatively correlated with recorded mean burst frequency, suggesting a relationship between extent of mechanosensitivity and bursting (Spearman’s rho = -0.62, p = 0.02).

### Bursts can synchronize widely along the spinal cord after SCI

Ongoing regular bursting was a predominant feature after SCI, where burst synchrony across roots was increased **(Figure 2A)**. Simultaneous synchronous bursts were common within ipsilateral DRs **(Figure 2B**_**1**,**2**_) while correlated contralateral activity showed both lag and lead times **(Figure 2C**_**1**,**2**_). To quantify the relationship between roots, cross-correlations were calculated for recordings with synchronous bursting **(Figure 2B**_**3**_, **2C**_**3**_). For the entire SCI population, ipsilateral bursts synchronized with limited lag, suggesting broad recruitment initiated by a common source **(Figure 2D)**. Contralateral bursting networks showed bidirectional interactions supporting capacity for coupling between distinct bursting networks **(Figure 2B)**. Overall, observed temporal relationships demonstrated coupling between discrete and overlapping burst generating networks.

**Figure 2.**
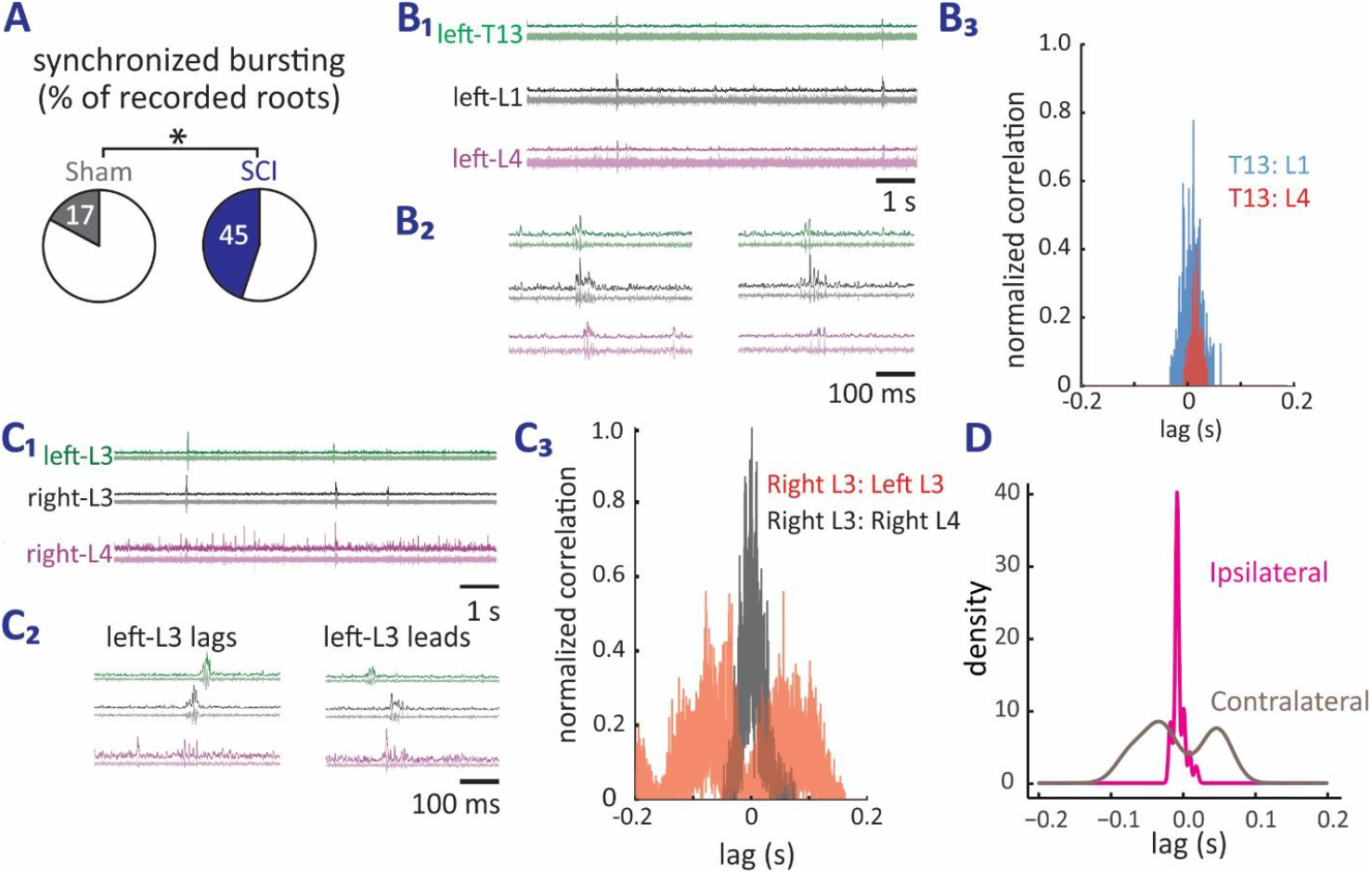
Coordination of bursting across segmental dorsal roots after SCI. **(A)** SCI preparations exhibited a significantly higher percentage of roots with synchronous bursting than did sham preparations (Chi-square (df=1, n = 13; 10 SCI, 3 sham) = 4.8, p-value = 0.03). **(B**_**1**_) Example bursting activity in three ipsilateral dorsal roots from another preparation. Two roots are adjacent (T13 and L1) while the other root is several segments caudal at L4. **(B**_**2**_) Magnified timescale examples. **(B**_**3**_)) Cross-correlograms visualizing 300 seconds of recordings reduced to only identified bursts with all background noise removed. Cross-correlograms between T13-L1 (shown in blue) and T13-L4 (shown in red) illustrate that bursts arrive almost simultaneously in all roots, with rostral T13 slightly leading the caudal L4 root. **(C**_**1**_) Example recording of bursting activity in three dorsal roots – two adjacent ipsilateral roots and one contralateral lumbar (L) root. Both raw and RMS amplitude filtered waveforms are shown. **(C**_**2**_) Magnified timescale examples of differing relationships between bursts shown above. In both cases, right L3 and right L4 occur near-simultaneously while contralateral left either L3 lags (left panel) or leads (right panel) bursting on right side. **(C**_**3**_) Cross-correlograms visualizing 300 seconds of recordings reduced to only identified bursts with all background noise removed. Right L3 and right L4 (shown in black) and right L3 and left L3 (shown in red) illustrate that the differences exemplified in B_1,2_ are consistently present throughout the entire recording. **(D)** Density plot (smoothed histogram) from cross-correlations across all SCI preparations reveals that coordinated bursts occur concurrently when roots were ipsilateral (n=8 mice) but with comparable lead/lag values when sampled roots were contralateral (obtained from the top 5 inter-root lag values; n=3 mice). Cross-correlations were calculated for all permutations of SCI roots with bursting in multiple roots.

### Bursting circuits can also be recruited by afferent stimuli and exhibit a refractory period

When spontaneous bursting was seen in any preparation, high-intensity afferent electrical stimuli (recruiting Aβ, Aδ and C fibers) also evoked bursts with comparable appearance to spontaneous bursts (n = 11/11 preparations) (**Figure 3A**_**1**_). To probe burst refractory state, we compared bidirectional interactions between spontaneous and evoked bursts. Afferent stimuli almost always failed to evoke a burst if a spontaneous burst occurred sooner than ∼700 ms before stimulation **(Figure 3A**_**2**_). Similarly, SCI spontaneous bursts following stimulus-evoked bursts almost never occurred sooner than ∼500 ms after stimulation **(Figure 3A**_**2**_). Overall, bursting events were also recruited by afferent stimulation, and timing interactions with spontaneous bursts revealed that recruitment was limited by a prolonged post-burst refractory period.

**Figure 3.**
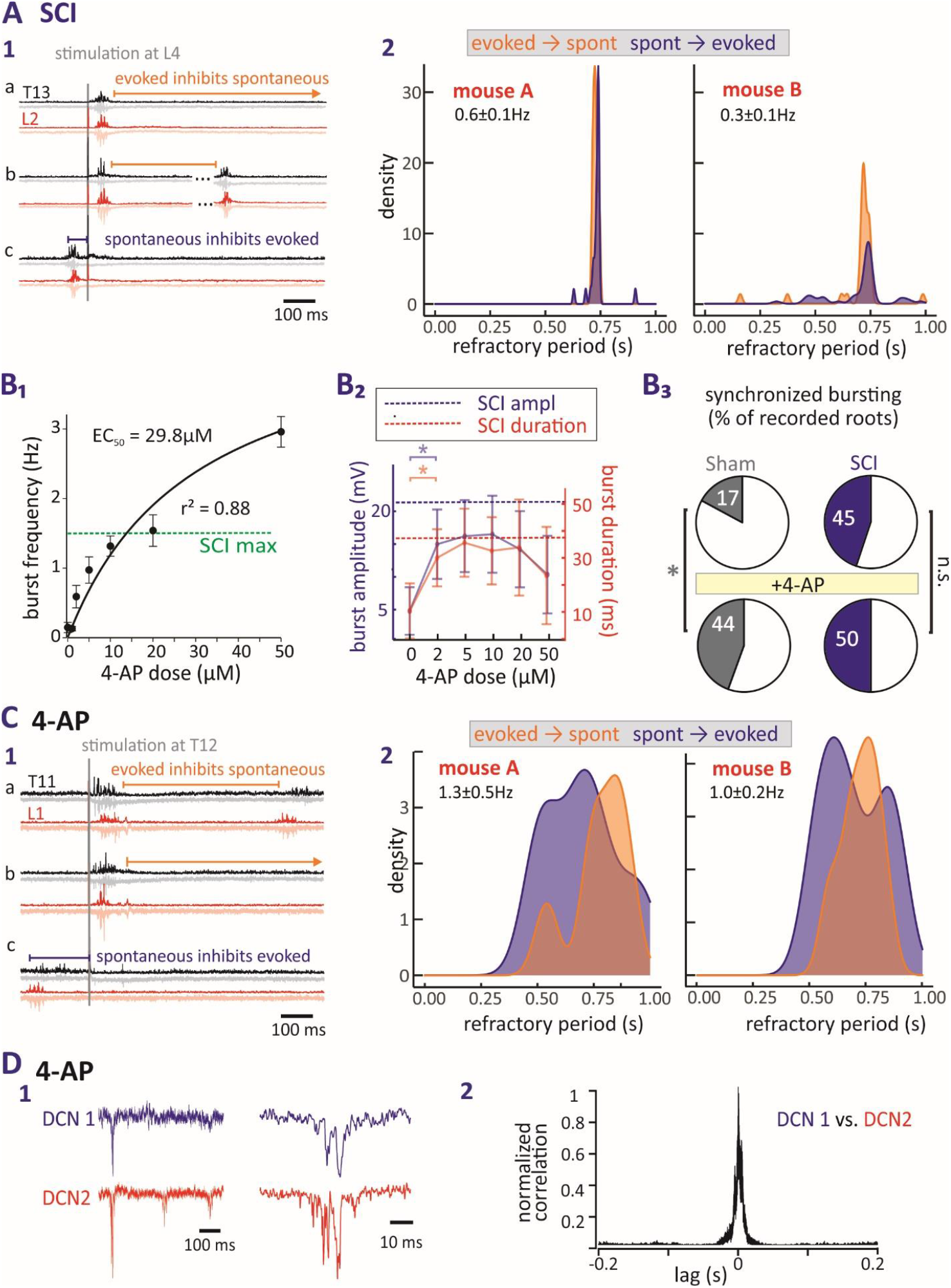
Relating bursting properties after SCI to those generated with 4-AP. **(A)** Mutually inhibitory interactions between spontaneous and evoked bursts were used to identify burst refractory periods in SCI mice. **(A**_**1**_) Example recording showing that spontaneous events are not seen for several hundred milliseconds after an evoked burst (panels a, b), and spontaneous bursts similarly prevent subsequent expression of evoked bursts (panel c). **(A**_**2**_) Density plots (smoothed histograms) quantifying the bidirectional distribution of burst refractory periods between spontaneous (spont) and evoked bursts in two SCI mice with mean 0.6 and 0.3 Hz spontaneous burst frequencies. Burst circuit refractory period was prominent for several hundred ms (bursts plotted: mouse A evoked-to-spont = 111, spont-to-evoked = 289; mouse B evoked-to-spont = 87, spont-to-evoked = 189). **(B**_**1**_) Effect of 4-AP dose on emergent burst frequency. Values shown are means ± standard error. Fit r^2^ = 0.88, EC_50_ = 29.8 μM (100 μM dose with value of 3.8± 0.2 Hz not shown). Dotted green line identifies SCI mouse with highest recorded burst frequency, which compares to frequencies obtained ∼20μM 4-AP (n=6). **(B**_**2**_) Burst amplitude and duration values were comparable at 4-AP doses between 2-20μM. Dotted lines representing mean amplitude and duration values from the SCI population show comparable values. (Welch’s t-test *p < 0.05, n=6) **(B**_**3**_) 4-AP (10μM) significantly increased the percentage of synchronously bursting roots in sham (Chi-square; p=0.02; n=7), but not SCI preparations (Chi-square; p=0.4; n=10). **(C)** Mutually inhibitory interactions between spontaneous and evoked bursts identify burst refractory period after 4-AP in naïve mice. **(C**_**1**_) Example recording showing that spontaneous events are not seen for several hundred milliseconds after an evoked burst (panels a, b), and spontaneous bursts similarly prevent subsequent expression of evoked bursts (panel c). **(C**_**2**_) Density plots quantifying the bidirectional distribution of burst refractory periods between spontaneous and evoked bursts in two mice undergoing 4-AP (10μM) induced bursting with mean 1.3 and 1.0 spontaneous burst frequencies. Note that refractory period duration was shorter than in SCI mice (possibly related to higher burst frequencies) (bursts plotted: mouse A evoked-to-spont = 127, spont-to-evoked = 296; mouse B evoked-to-spont = 53, spont-to-evoked = 292). **(D)** 4-AP induced antidromically propagating synchronous bursting in adjacent dorsal cutaneous nerves (DCN) *in vivo* (dose; 0.2 mg/kg). **(D**_**1**_) Raw recording examples of temporally associated bursts in adjacent DCNs. **(D**_**2**_) Cross-correlogram indicates that DCN bursts are synchronous (total number of bursts plotted = 811).

### The convulsant 4-aminopyridine generates bursting properties comparable to that seen after SCI

As synchrony and refractory periods are features of epileptiform activity, we compared SCI bursting with that induced by 4-aminopyridine **(4-AP)**, a convulsant used in animal models of epilepsy [51] and known to generate epileptiform activity in DH neurons *in vitro* [40] and synchronous bursting in primary afferents *in vivo* [52]. 4-AP recruited spontaneous bursting activity in all naïve preparations (n=6/6) with a dose-dependent effect on burst frequency **(Figure 3B**_**1**_). Above a small dose of 4-AP, burst amplitude and duration plateaued at a level comparable to that seen in SCI mice **(Figure 3B**_**2**_). Strikingly, the number of roots with synchronized bursting dramatically increased after 4-AP administration in sham but not in SCI preparations **(Figure 3B**_**3**_), suggesting that 4-AP and SCI recruit a common bursting circuitry. Spontaneous burst waveform, inter-burst interval, and recruitment by afferent stimulation were also comparable between 4-AP and SCI preparations, suggesting hyperexcitability recruited similar circuits in the two conditions **(Figure 3C**_**1**_**-3C**_**2**_).

To confirm that synchronous bursting occurred at physiological temperature and propagated peripherally, separate *in vivo* experiments were undertaken while recording from adjacent dorsal cutaneous nerves. Administration of 4-AP induced synchronized bursting recorded in dorsal cutaneous nerves (n=2/2) **(Figure 3D**_**1-2**_), similar to that previously reported in the acutely spinalized cat [53].

### GABA_A_-receptor mediated actions drive emergence of ectopic busting

Normally, dorsal root stimulation evokes primary afferent depolarization – recorded as subthreshold DRP responses - as a form of negative feedback presynaptic inhibition [13, 14]. Evoked and largely subthreshold DRPs were seen with conversion to excitatory bursting after 4-AP, and evoked actions were subsequently greatly depressed or blocked with the GABA_A_ receptor antagonist bicuculline (n=4/4; **Figure 4A**). In all mice, 4-AP initiated bursts or increased spontaneous burst frequency, and bursts were always blocked by GABA_A_ receptor antagonists (SCI n=8; naïve n=2; and sham n=7) **(Figure 4B)**. GABA_A_ block of spontaneous bursting was also seen in 2/2 SCI preparations never given 4-AP **(Figure 4C)**. That both SCI- and 4-AP-induced ectopic bursting required GABA_A_ receptors provides additional support that bursts are driven by common interneurons and further highlights the similarity between SCI and 4-AP-evoked bursting.

**Figure 4.**
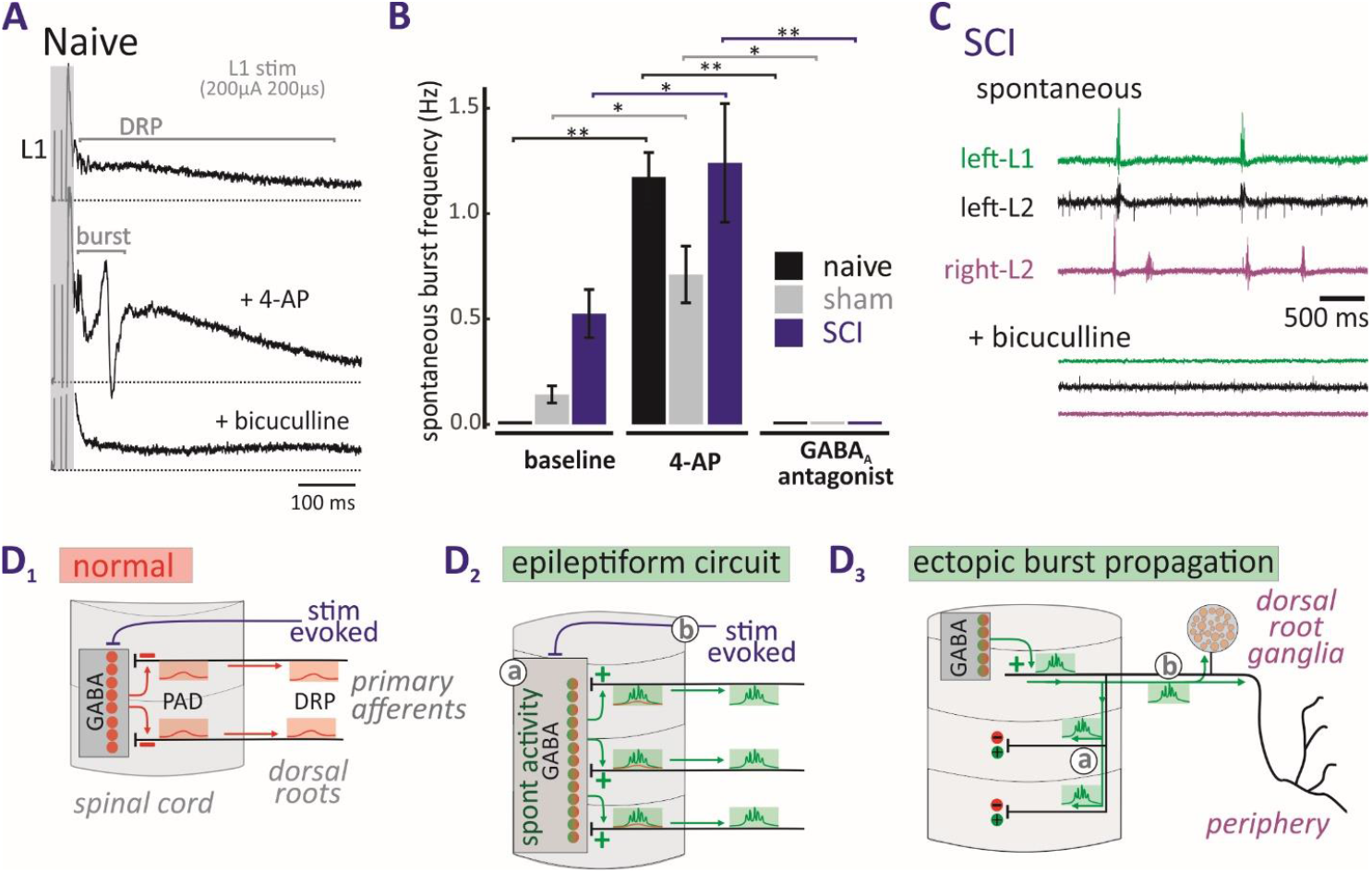
Role of GABA_A_ receptors in the generation of ectopic bursting. **(A)** Example of DR stimulation-evoked subthreshold dorsal root potential (DRP) in a naïve preparation (top). After addition of 10μM 4-AP, DRP is increased in amplitude and a suprathreshold volley is also present (middle). Evoked bursting and DRP are blocked by bicuculline (bottom) (n = 4). DRP and bursting are identified with horizontal bars. Dotted lines indicate baseline. Traces are averages of 10 events. **(B)** Effect of 4-AP and bicuculline on spontaneous burst frequency in naïve (n=2), sham (n=2-7 per drug condition), and SCI (n=6-10 per drug condition) preparations. Spontaneous bursting is induced or increased in frequency by 4-AP (10μM). Bursting is subsequently completely blocked in similar fashion with GABA_A_ antagonists bicuculline (n = 5), picrotoxin(n=3), and gabazine(n=4) so results are pooled. (Welch’s t-test *p < 0.05, **p<0.01, ***p<0.001; n=2-10 depending on drug and injury condition; values show mean ± standard deviation). **(C)** Spontaneous bursting in SCI cord blocked with bicuculline. **(D)** Outcome of hyperexcitability induced bursting after SCI. Panels **D**_**1 &**_ **D**_**2**_ show afferent input onto intraspinal GABAergic interneurons and their axoaxonic synaptic actions on intraspinal afferents. **(D**_**1**_) Normally, afferent input acts on interposed GABA interneurons to provide negative feedback presynaptic inhibition via subthreshold afferent membrane depolarization (PAD) recorded as dorsal root potential (DRP) (orange boxes). **(D**_**2**_) SCI or 4-AP supports multisegmental synchronization of GABAergic PAD circuits. (**a**) Emergent spontaneous synchronized GABAergic drive increases PAD leading to suprathreshold ectopic bursts that propagate across multiple spinal segments (green boxes). (**b**) Afferent stimulation can recruit the same circuitry. **(D**_**3**_) Ectopic bursting propagates bidirectionally. (**a**) Reentrant propagation along afferent collateral projections lead to activation of spinal circuits. (**b**) Antidromically-propagating bursts project to cell bodies in dorsal root ganglia and peripheral terminal fields. Additional studies are required to identify contributing populations of interneurons, mechanisms of circuit synchronization, the identity of afferents that recruit and are recruited by bursting circuits, and their contribution to sensory dysfunction after SCI.

## Discussion

These results demonstrate that SCI promotes the emergence of epileptiform circuit behavior with GABAergic neurons implicated as drivers of aberrant ectopic bursting in intraspinal afferents. Observed epileptiform characteristics included episodes of stereotyped bursting activity, hypersynchrony, burst triggering by sensory input, post-burst refractory period, and fundamental changes in circuit behavior.

A previous group focusing on hamster found that the isolated *ex-vivo* spinal cord develops spontaneous afferent bursting over time [27, 54]. Reexamination of naïve mice in an earlier study in isolated cord [50] showed similar time-dependent development of spontaneous activity in 3/24 mice, demonstrating that the DH circuitry driving ectopic bursting is normally suppressed. As spinalization facilitates synchrony of GABAergic interneuron actions on afferents [26], it is likely that SCI induced loss of descending modulatory tone enables the emergence of synchronous epileptiform bursting. 4-AP also recruited epileptiform ectopic bursting in primary afferents. The ability to recapitulate SCI-induced epileptiform bursting with this K^+^ channel blocker is consistent with known reductions in voltage-gated K^+^ channel function observed after SCI [30, 32] and with SCI-induced conversion of GABA interneurons into a bursting phenotype [55].

The generation of larger DRPs due to stronger GABAergic depolarization is likely responsible for conversion of subthreshold DRPs to suprathreshold bursts **(Figure 4A)** as depolarization toward E_GABA_ in afferents, thought to be ∼ -30mV [17], should be above spike threshold. This conversion [13, 56] represents a fundamental functional perturbation in sensory system control, as it indicates that subthreshold negative feedback (**Figure 4D**_**1**_) now also includes aberrant suprathreshold ectopic spiking (**Figure 4D**_**2**_). Spiking is expected to influence system excitability via reentrant central collaterals and peripheral projections (**Figure 4D**_**3**_). Reentrant excitatory synaptic actions may act on multiple pathways but are unlikely to recruit the same GABAergic presynaptic circuit due to their arrival during the epileptiform refractory period.

Overall, these results support a conceptually novel understanding of SCI -induced sensory dysfunction. The observed epileptiform phenotype of segmentally propagating bursts is consistent with descriptions of propagating neuropathic pain (e.g. shooting, radiating, stabbing)[1] with common pharmacologic control [40] and hence provides a novel hypothetical substrate to explore underlying mechanisms and their therapeutic control.

## Materials and methods

### Animals

All procedures were approved by the Emory University Institutional Animal Care and Use Committee. The primary cohort of mice were SCI (n=10) and sham (n=7) littermates from 3 litters aged between 110 and 130 days at time of surgery, and between 310 and 640 postnatal days at time of terminal experiments. Post-surgery, animals were housed in isolated cages with a maximum of two animals per cage. The wide range of ages at terminal experimentation is due to delays associated with the COVID19 pandemic. Similar behavioral and electrophysiological phenotypes have been confirmed in ongoing studies in mice aged P60 at surgery and ∼P100 at sacrifice. Naïve mice used for comparison to sham/SCI cohort were a male P117 and a female P180. Naïve mice used for 4-AP concentration experiments were 4 male and 2 female aged between 241 and 248; naïve mice assessed for spontaneous bursting were 14 male and 8 female aged between 90 and 150 days at terminal experimentation.

### Spinal Cord Injury

Contusion injuries were performed as previously described using the Infinite Horizon spinal cord impactor device [57]. Briefly, mice were anesthetized with 5% inhaled isoflurane, then a midline incision and dorsal laminectomy performed at T10-T12 based on distance from cord apex. The mouse was moved to the impactor and the exposed cord impacted at 50kD with 0s dwell time. Mice were administered 2 mg/kg meloxicam (Cayman Chemical Company) the day of surgery and the next day, as well as 0.5 mg/kg Enrofoxacin (Baytril, Bayer) daily following surgery for 7 days. Sham mice underwent identical surgical and post-surgical procedures other than cord contusion.

### Measurement of paw withdrawal reflex changes

Mice underwent von Frey testing for 2 weeks before injury, as well as after injury to confirm development of mechanical hypersensitivity as a proxy for allodynia. Mechanical hypersensitivity was measured using Chaplan’s up-down protocol for von Frey filaments [58]. Briefly, animals were acclimated to the von Frey filaments (0.4, 0.6, 1.4, and 2.0 grams), cages, and mesh floor for several days prior to testing. Mechanical sensitivity was assessed in both hind paws three times prior to surgery and once weekly afterwards for 10 weeks. The 50% paw withdrawal threshold (**PWT**) was quantified for SCI and sham groups. This metric is defined as the stimulus intensity (in grams) required to produce a withdrawal response 50% of the time and is a common measure of mechanical allodynia in both human and animal populations [59, 60].

### Dissection for *ex vivo* spinal cord experimentation

Mice were lightly anesthetized with inhaled isoflurane then injected intraperitoneally with 100μL 50% urethane for deep anesthesia. To induce hypothermia, dorsal skin overlying the vertebral column was removed and mice were submerged in ice-cold artificial cerebrospinal fluid (**aCSF**) until respiration rate slowed (2-3 minutes). Animals were then removed, decapitated, and the whole spinal cord dissected with a ventral approach as previously described [50]. The spinal cord dissection was performed in ice cold recovery aCSF [61] oxygenated with 95% O_2_-5%CO_2_. The isolated cord was then equilibrated to room temperature for 1 h in modified HEPES holding solution [61] oxygenated with 95% O_2_-5%CO_2_, and then pinned dorsal side up in a Sylgaard-lined recording chamber while superfused with an oxygenated aCSF containing (in mM) 128 NaCl, 1.9 KCl, 1.3 MgSO4, 2.4 CaCl2, 1.2 KH2PO4, 10 glucose, and 26 NaHCO3 at ∼40 ml/min. All experiments were undertaken at room temperature.

### Dissection for *in vivo* dorsal cutaneous nerve recordings

Using a SomnoSuite (Kent Scientific) anesthesia machine, deep anesthesia was induced with 4% inhaled isoflurane and maintained with 1-3% isoflurane based on animal respiration and heart rate. Body temperature was maintained with a disposable hand warmer. 0.2 mg/kg 4-AP was administered intraperitoneally once deep anesthesia was established and the animal showed no response to toe pinch with forceps. A dorsal midline incision was performed, and dorsal cutaneous nerves freed along one side of the animal to allow for recording of adjacent nerves. A small amount of aCSF was pooled around the nerves to allow recording.

### Electrophysiology

Suction electrodes were fabricated from 1.65/0.75 (OD/ID) glass capillary tubes (Dagan Corp) using Narishige PC-100 electrode puller with tips broken back to achieve internal tip diameters of 100-250 μm. Electrodes were placed on various dorsal root entry zones for recordings and at more distal sites for recording or stimulation. Electrical stimulation was delivered using constant current stimuli [62] to the distal ends of dorsal roots (Fig 1B). All recorded data were digitized at 10 kHz (Digidata 1322A 16 Bit DAQ, Molecular Devices, U.S.A.) with Clampex acquisition software (v. 10.7 Molecular Devices). Recorded signals were amplified (10000x) and low-pass filtered at 3 kHz using in-house amplifiers. In all data presented, the number of animals utilized for analysis is represented by the noted n value. For each animal, a representative value was determined by averaging the evoked response from multiple trials within that animal (a minimum of 4) or by analyzing a minimum of 5 minutes of spontaneous gap-free recording.

### Drug administration

All drugs were applied to 100 mL recirculating bath. The convulsant 4-aminopyridine (**4-AP**, 1-100 μM, Spectrum) [30] was used to generate a model of sensory circuit hyperexcitability in the *ex vivo* intact spinal cord preparation and *in-vivo* (0.2mg/kg) to confirm peripheral propagation at physiological temperature. 4-AP has been demonstrated to increase the excitability of neurons in preclinical and clinical studies [63, 64] and recruits spinal nociception-encoding circuitry consistent with the emergence of spontaneous neuropathic pain [40, 65]. To characterize the dose-response relationship of the model, 4-AP was bath applied at increasing concentrations during recording of dorsal roots. To assess contributions from GABA_A_ receptors, the following antagonists were tested: bicuculline (10 μM, Enzo Life Sciences), gabazine (1 μM, EMD Millipore), and picrotoxin (25 μM, Tocris Bioscience). The NMDA antagonist APV (20μM, Tocris Bioscience) and AMPA/kainate antagonist CNQX (10 μM, Tocris Bioscience) were used to block ionotropic glutamatergic synaptic activity.

### Data and statistical analysis

Captured data was analyzed in pClamp, Spike2 (Cambridge Electronic Design), and custom-written scripts and applications in Python and R (see code accessibility section for access to Python analysis program). Recorded roots were selected for analysis based on the presence of bursting at baseline or assumption of viability due to observed occasional spikes or after addition of 4-AP. To assess below-level changes in somatosensory circuitry, only roots T13-L4 were considered for analysis. Any roots that did not show bursting after 4-AP addition were assumed damaged from dissection and excluded from analysis. Roots with signal to noise ratios too poor to reliably differentiate bursts from background noise were also excluded from analysis. Post-burst refractory period analysis involving stimulation, all electrical stimulus strengths that resulted in a burst (not direct afferent volley) were considered as a single population. Statistical significance of data was determined Spearman’s correlation, Chi-square, Wilcoxon rank-sum test when appropriate (as determined by Shapiro-Wilks test), or Welch’s two-tailed t-test, depending on the normality and scedasticity of the data. Relationships were considered significant at p < 0.05. See figure legends for individual statistical tests and outcomes. Values are expressed as mean ± standard deviation unless otherwise stated.

Spontaneous and evoked afferent bursts were identified and quantified with custom-written Python software. Briefly, peaks were identified with a peak threshold set at 3.5x RMS noise. Recordings were considered to contain bursting activity when the standard deviation of inter-spike intervals was greater than the mean inter-spike interval of all peaks [66]. Inter-spike interval was also used to characterize packets of peaks as distinct individual bursts (peaks were binned together when they were <20 ms apart and only groups of 5 or more peaks were considered a burst). Once bursts were identified, they were quantified (amplitude, duration, etc.) and cross-correlations were processed with all possible permutations of channels taken 2 at a time. Data from this program were exported and analyzed and visualized with R.

## Code accessibility

Custom-written Python application for burst analysis is available at https://github.com/mbryso4/2023_Burst_Analysis. Available versions are the same that were used for analysis of this work. Updated versions will be maintained elsewhere.

## Acknowledgments

Supported by NIH NS102850 and PVA Foundation. We thank Dr. Peter Wenner for reading an earlier version of the manuscript.

